# Behavioral and brain mechanisms mediating conditioned flight behavior in rats

**DOI:** 10.1101/2021.01.29.428848

**Authors:** Michael S. Totty, Naomi Warren, Isabella Huddleston, Karthik R. Ramanathan, Reed L. Ressler, Cecily R. Oleksiak, Stephen Maren

**Author notes:** <u>Corresponding author</u>*, Address: Stephen Maren, Department of Psychological and Brain Sciences Texas A&M University, 301 Old Main Dr., College Station, TX 77843-3474.

## Abstract

Environmental contexts and associative learning can inform animals of potential threats, though it is currently unknown how contexts bias defensive transitions. Here we investigated context-dependent flight responses in the Pavlovian serial-compound stimulus (SCS) paradigm. We show here that SCS-evoked flight behavior in male and female rats is dependent on contextual fear. Flight was reduced in the conditioning context after context extinction and could be evoked in a different shock-associated context. Although flight was exclusive to white noise stimuli, it was nonetheless associative insofar as rats that received an equal number of unpaired USs did not show flight-like behavior. Finally, we found that inactivation of either the central nucleus of the amygdala (CeA) or bed nucleus of the stria terminalis (BNST) attenuated both contextual fear and flight responses. This work demonstrates that contextual fear summates with cued and innate fear to drive a high fear state and freeze-to-flight transitions.

## INTRODUCTION

The selection of appropriate defensive behavior is vital to survival in the face of threat. Associative learning allows animals and humans to adapt their behavior to avoid predicted danger, and environmental contexts are critical for discriminating between fear and safety. Traumatic events can lead to pathological fear and the dysregulation of contextual processing appears to be central to various psychopathologies, such as post-traumatic stress disorder (PTSD) (Liberzon and Abelson, 2016; Liberzon and Sripada, 2008; Maren et al., 2013). For example, context processing deficits in patients with PTSD can lead to the overgeneralization fear (Grillon and Morgan, 1999; Jovanovic et al., 2012; Kaczkurkin et al., 2017; Lis et al., 2020; Morey et al., 2020, 2015; Orr et al., 2000), deficits in the extinction of fear (Blechert et al., 2007; Jovanovic et al., 2012; Milad et al., 2009; Norrholm et al., 2011; Rougemont-Bücking et al., 2011; Steiger et al., 2015; Wicking et al., 2016), and the renewal of extinguished fear in safe contexts (Garfinkel et al., 2014). This suggests that a complete understanding of how contexts regulate conditioned defensive behavior is essential to identifying neural circuits relevant to fear and anxiety disorders.

Pavlovian fear conditioning has been used for decades to model aversive learning and memory in rodents. Previous work has revealed that contexts can both directly elicit conditioned fear responses and also act as occasion setters to gate the retrieval of aversive memories (Bouton, 2002; Fraser and Holland, 2019; Maren et al., 2013). In both cases, fear responses in rodents typically manifest as defensive freezing behavior, which is used as the primary metric of conditioned fear in rodents. Predatory imminence theory posits that defensive behavior scales with threat proximity on a spatiotemporal scale such that freezing behavior is seen in post-encounter modes (once threat has been realized) whereas flight behavior is part of the circa-strike defensive mode (when threat is proximal) (Fanselow and Lester, 1988). This has been demonstrated in both humans and rodents using naturalistic predator threats (Mobbs et al., 2007; Yilmaz and Meister, 2013), but it remains unclear whether conditioned threats (such as auditory conditioned stimuli) can drive circa-strike behavior such as flight. Recently, Fadok and colleagues developed a modified auditory Pavlovian fear conditioning procedure that uses a serial-compound stimulus (SCS) to elicit both freezing and flight defensive modes in mice (Fadok et al., 2017). The SCS is comprised of a pure tone stimulus immediately followed by a white noise stimulus that elicits two conditioned responses (CRs): freezing and flight behavior, respectively. Moreover, they show that the switch between freezing and flight behavior is gated by microcircuitry within the central nucleus of the amygdala (CeA), a structure critical to the expression of Pavlovian CRs (Janak and Tye, 2015; Killcross et al., 1997). Interestingly, in the SCS paradigm flight behavior is normally limited to the conditioning chamber and freezing dominates when the SCS is presented in a different context. This procedure presents a unique opportunity to investigate mechanisms by which context and associative memory may scale between freezing and flight defensive modes.

In order to fully reveal the neural circuits that may underly pathological fear, we must understand the distinct neural circuits that underly various defensive modes and how they may be gated or modulated by context (Mobbs et al., 2020). Here we sought to determine the behavioral and neural mechanisms mediating the influence of context on the expression of defensive behaviors to an SCS. One possibility is that context serves as an occasion setter and promotes SCS-evoked flight behavior in the conditioning context, but not in other contexts. Another possibility is that direct context-US associations produce fear that summates with that to the SCS to elevate threat imminence thereby yielding flight. The occasion setting hypothesis predicts that flight would be specific to the conditioning context and would not be expressed elsewhere, whereas the summation hypothesis predicts that flight would be evoked in any shock-associated context, regardless of whether it had hosted SCS-shock trials. In a series of experiments to test these competing hypotheses, we found that rats displayed flight behavior when the SCS was presented in a US-associated context different than the conditioning context. Moreover, extinguishing fear to the conditioning context suppressed flight behavior in that context. We further provide evidence that SCS-evoked flight is a conditioned response by showing that flight-like behavior cannot be explained by sensitization or fear-potentiated startle. Finally, we show that pharmacologically inactivating either the CeA or BNST, brain regions that are critical to the expression of contextual fear, reduces flight-like behavior. We thus argue that SCS-evoked flight behavior is a high fear state driven by the summation of cued, contextual, and innate fear.

## RESULTS

### A conditioned serial-compound stimulus evokes flight behaviors in rats

Previous work shows that SCSs can evoke flight behavior in mice, but it is unknown if this behavior occurs in rats. Therefore, we first sought to determine if rats show flight-like behavior to an SCS using the behavioral protocol first described by Fadok and colleagues (Fadok et al., 2017). In this procedure (**Figure 1A**), rats were first habituated to four SCS presentations (tone→white noise; each stimulus consisted of 10 sec trains of 500 ms pips with an inter-pip interval of 500 ms) in context A (Day 1), then conditioned with five SCS-US presentations for the next three days in context B (Days 2-4), and finally tested with four SCS-alone presentations in both context A and B (Days 5 and 6; counterbalanced). For this experiment, we quantified 1) freezing as a percentage of time, 2) average motor activity, as well as 3) the number of jumps (all four paws leaving the floor) and darts (rapid movement from one position to another) during both tone and noise components of the SCS. Freezing and activity were quantified automatically online by digitizing voltages emitted by force transducers under each chamber; jumps and darts were scored offline from video recordings of the sessions by observer’s blind to the experimental conditions.

**Figure 1.**
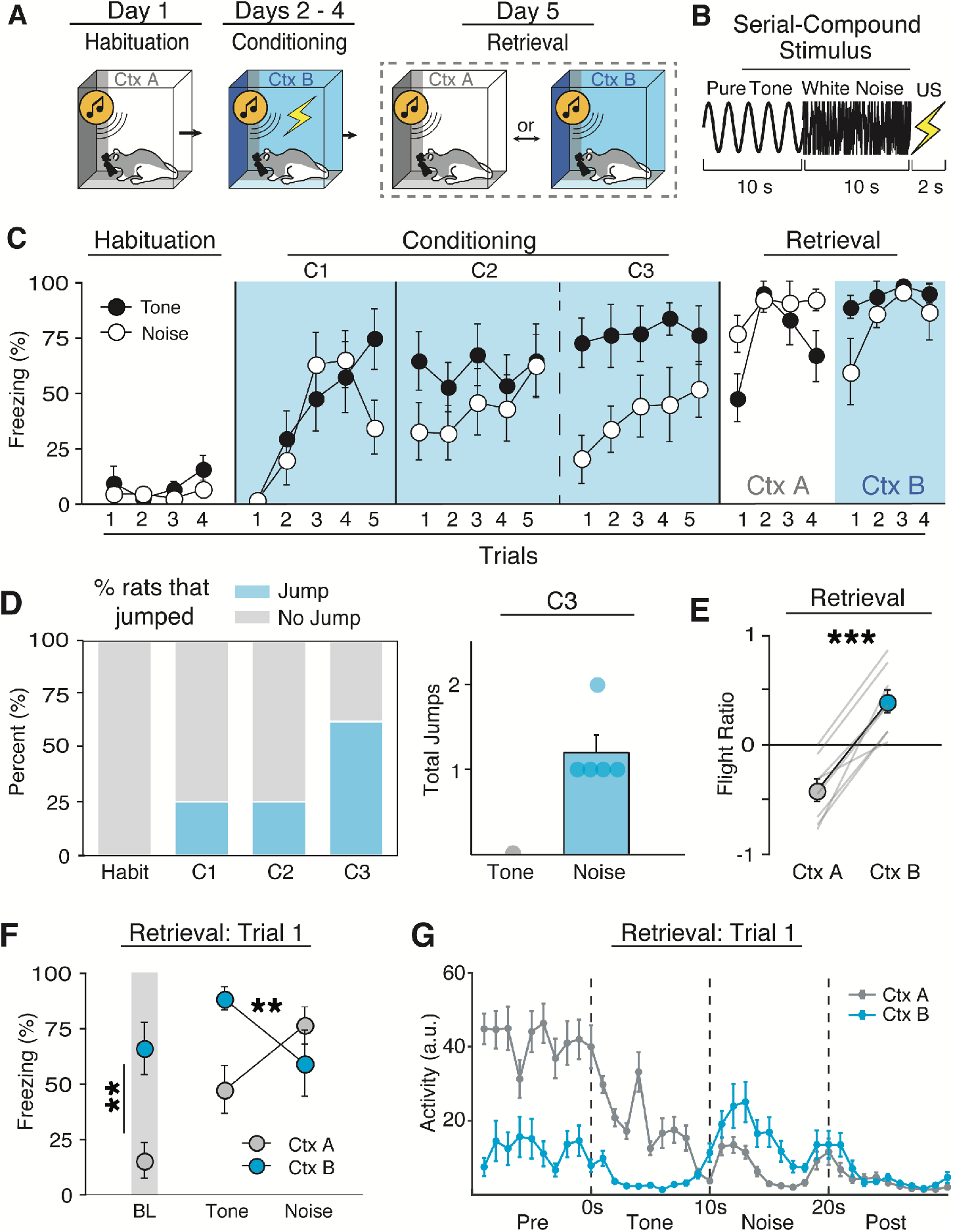
Rats display context-dependent flight-like behavior to a conditioned SCS. **A**, Schematic of the behavioral design used in Experiment 1. **B**, Schematic representation of the serial-compound stimulus (SCS). **C**, Average freezing data for tone and noise stimuli during each SCS presentation during Habituation, Conditioning, and Retrieval. Rats showed lower freezing to the noise on the second and third day of conditioning. **D**, Percentages of rats that showed at least one jump during an SCS for each respective day of behavioral testing. Of the rats that showed at least one jump, jumps were exclusive to noise stimuli (C3). **E**, Average flight ratio in which positive numbers represent increased movement to the noise relative to tone, whereas negative numbers represent decrease activity relative to tone. Rats displayed flight like behavior when tested in the conditioning, but not habituation context. **F**, Average freezing data during Retrieval shows that rats tested in the conditioning context showed high freezing during baseline and the first tone presentation but decrease to the noise, whereas rats tested in the habituation context showed low freezing to baseline and the tone which increase to the noise. **G**, Averaged motor activity data from 10s before SCS onset to 10s after SCS offset in both the Habituation (Ctx A) and Conditioning (Ctx B) contexts. All data are represented as mean ± SEM; *, **, and *** denotes *p*<.05, *p*<.01, and *p*<.001, respectively.

As shown in **Figure 1C**, prior to conditioning, SCS presentations produced low levels of freezing and there was no difference in stimulus type on either freezing [*F*(1, 6) = 1.31, *p* = .295] or activity [*F*(1, 6) = 2.57, *p* = .160]. All rats showed increased freezing behavior throughout the first day of conditioning [main effect of trial, *F*(4, 24) = 8.94, *p* < .0001]. Although there was no main effect of stimulus type [*F*(1, 6) = 1.91, *p* = 2.16], there was a trial x stimulus type interaction [*F*(4, 24) = 3.33, *p* = .026] with noise producing a decrease in freezing relative to the tone stimulus on the last trial of the first conditioning session (**Figure 1C**). This suggested that noise onset was associated with a reduction in freezing. Indeed, on the second day of conditioning rats showed less freezing to the noise CS [*F*(1, 6) = 10.12, *p* = .019], which was mirrored by an increase in activity [*F*(1, 6) = 15.27, *p* = .008]. Although the rats displayed a clear switch in defensive behavior upon noise onset, the number of jumps to the noise were low with only ~25% of rats displaying at least one jump (**Figure 1D**). During this third and final conditioning session, all rats showed an even greater decrease in freezing and increase in activity upon noise onset [Freezing: main effect of stimulus type, *F*(1, 6) = 52.23, *p* = .0004; Activity: main effect of stimulus type, *F*(1, 24) = 41.15, *p* = .001], and ~60% of the rats performed at least one jump. Importantly, jumps were nearly exclusive to noise presentations, with only three total jumps observed during tone presentations across all three days of conditioning. Despite a clear increase in activity, rats emitted only a small number of total jumps with only one rat jumping multiple times during the third conditioning session (**Figure 1D**).

Recent reports have noted that aversive CSs can elicit darting behavior, particularly in female rats (Gruene et al., 2015). However, we seldomly observed darting behavior to the SCS in male or female rats (3 or less total darts across all animals each day). Moreover, there were no sex differences in freezing [main effect: *F*(1, 6) = .769, *p* = .414] or activity [main effect: *F*(1, 6) = .598, *p* = .469] evoked by the SCS across conditioning. In summary, a conditioned SCS elicits a clear switch from freezing to activity with infrequent jumps in both male and female rats. It appears that an increase in motor activity (and decrease in freezing) is the dominant mode of SCS-evoked flight-like behavior in rats, compared to frequent jumping previously observed in mice (Fadok et al., 2017; Hersman et al., 2020). We will therefore use white noise-evoked decreases in freezing from here on as the primary metric for flight behavior in rats.

### SCS-evoked flight behavior in rats is context-dependent

Next, we sought to determine if flight behavior in rats is context-dependent as previously shown in mice (Fadok et al., 2017). In previous work, flight behavior was observed in the conditioning context, but not when the SCS was presented in the habituation context. To test this, conditioned rats were placed into either the habituation or conditioning context (Ctx A and B, respectively) and presented the SCS four times without the US. Although there was no difference in overall freezing between contexts [*F*(1, 6) = .95, *p* = .367] or stimulus type [*F*(1, 6) = .14, *p* = .720], rats displayed a clear decrease in freezing upon noise onset in the conditioning context (**Figure 1C & F**), whereas they instead froze more to the noise presentation in the habituation context [context X stimulus type interaction, *F*(1, 6) = 17.82, *p* = .006]. This was again mirrored by activity levels [*F*(1, 6) = 14.12, *p* = .009]. In other words, rats displayed an increase in activity upon noise onset in the conditioning context, but not the habituation context (**Figure 1G**). There was no main effect of sex for either freezing [*F*(1, 6) = .434, *p* = .535] or activity levels [*F*(1, 6) = .770, *p* = .414]. Interestingly, flight only occurred during the first trial of retrieval testing and rats froze at high levels for the remainder of the test trials [main effect of trial, *F*(3, 18) = 8.25, *p* = .001]. This is reminiscent of the rapid extinction of flight behavior previously reported in mice (Fadok et al., 2017).

**Figure 1G** shows that when tested in the conditioning context rats exhibited low levels of activity to the tone but increased their activity upon white noise onset. However, in the habituation context, rats exhibited low levels of activity throughout the duration of the SCS. To further quantify this, we computed a “flight ratio”, which was the ratio of the difference of noise and tone load-cell activity to the sum of noise and tone load-cell activity for the first retrieval test trial (further described in the methods). This metric spans a scale from −1 to 1 whereby increased activity during noise relative to tone is represented as positive values and decreased relative activity is represented as negative values. As shown in **Figure 1E**, flight ratios were greater to the noise compared to the tone in the conditioning context relative to the habituation context [*F*(1, 6) = 86.26, *p* < .0001]. There was once again no main effect of sex [*F*(1, 6) = .041, *p* = .911]. This shows that the SCS-driven flight behavior observed in rats is limited to the conditioning context. Collectively, these results demonstrate that a conditioned SCS drives flight-like behavior in rats manifest as a reduction in freezing punctuated by infrequent jumping behavior, and this pattern of responding to the SCS was context-dependent, as has previously been reported in mice.

### Flight-like behavior depends on context-US associations

We next investigated what properties of the test context gates flight behavior. One possibility is that context serves as an occasion setter, informing the animal about the SCS-US association in the conditioning context. Alternatively, flight may be driven by a high fear state resulting from the summation of SCS-US and context-US associations. To discriminate among these possibilities, we explored whether conditioned flight would be expressed in an excitatory context that had never hosted SCS-US trials (i.e., a context in which animals experienced unsignaled shocks).

To this end, rats first underwent habituation and conditioning as previously described (**Figure 2A**). There was once again very little freezing to the SCS prior to conditioning and no difference in stimulus type [*F*(1, 12) = 1.607, *p* = .229]. Rats displayed increased freezing across conditioning sessions [main effect of day: *F*(2, 24) = 9.809, *p* = .0008] as well as noise-elicited decreases in freezing [day X stimulus type interaction: *F*(1, 12) = 14.109, *p* < .0001]. Next, to test if flight depends on a context-US association, rats were separated into two groups that would either receive five unsignaled USs in a novel context (context C) or would merely be exposed to the same context for an equal amount of time (**Figure 2A**, **Day 5**). As shown in **Figure 2B**, rats that received unsignaled USs (Shock group) showed increased freezing across the session, whereas rats that were not shocked (No-Shock group) froze at low levels [minutes X group interaction: *F*(5, 60) = 11.364, *p* < .0001].

**Figure 2.**
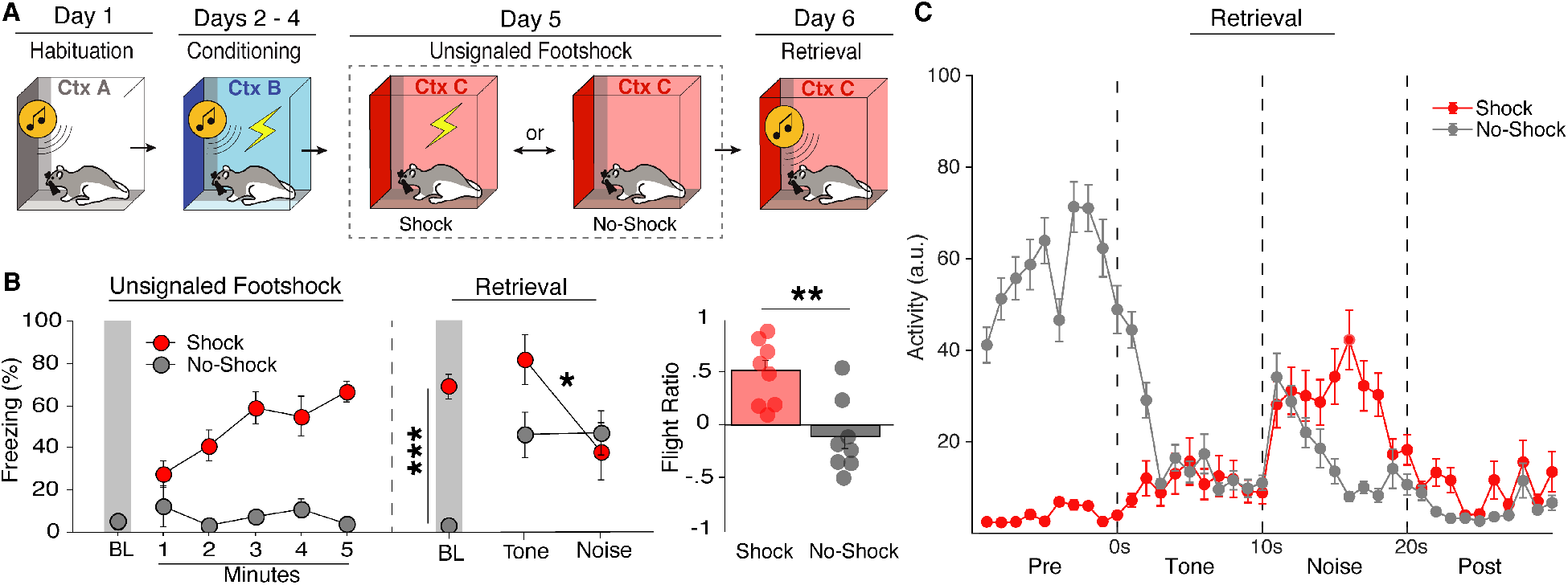
Flight-like behavior depends on context-US associations. **A**, Behavioral design for Experiment 2. **B**, Average freezing data shows rats that received unsignaled footshocks (Shock) froze at high levels, whereas rats that were merely exposed to the context (No-Shock) froze at low levels. For Retrieval testing, Shock animals showed higher baseline freezing and a decrease in freezing upon white noise onset whereas No-Shock animals showed low baseline levels and remained freezing at moderate levels throughout the SCS. **C**, Average flight ratio shows that rats that Shock animals showed a bigger flight response than No-Shock animals. **D**, Averaged activity data during the first trial of Retrieval for Shock and No-Shock animals. All data are represented as mean ± SEM; *, **, and *** denotes *p*<.05, *p*<.01, and *p*<.001, respectively.

For retrieval testing, all rats were placed back into context C and after a 3-minute baseline period were presented one SCS-alone trial. Shock animals showed much higher levels of fear to the context via freezing during the baseline period compared to No-Shock animals [*F*(1, 12) = 107.324, *p* < .0001]. Upon SCS presentation, Shock animals displayed a dramatic switch from freezing to activity upon noise onset (**Figure 2C**), whereas No-Shock animals decreased freezing momentarily, but quickly reverted back to freezing [group X stimulus type interaction: *F*(1, 14) = 5.928, *p* = .0289]. This was mirrored by the flight ratio [*F*(1, 12) = 12.926, *p* = .0037] (**Figure 2B**). In other words, conditioned animals presented the SCS in a shock-associated context displayed flight-like behavior, whereas animals tested in a neutral context did not. No sex differences were seen during retrieval [*F*(1, 12) = .498, *p* = .494] nor at any other point in this experiment. Thus, by showing that flight behavior can be evoked in a shock-associated context different from the original conditioning context, these results demonstrate SCS-driven flight-like behavior depends on context-US associations rather than occasion setting by the conditioning context. This suggests that flight to an SCS is driven by a high fear state gated by summation of SCS-US and context-US association.

### Extinguishing contextual fear reduces flight-like behavior

If contextual fear drives flight to an SCS, then extinguishing that fear should reduce flight behavior. To test this, we habituated and conditioned rats as previously described (**Figure 3A**), and then extinguished the conditioning context prior to retrieval testing. As in the previous experiments all rats showed similarly low levels of freezing to both stimuli prior to conditioning [*F*(1, 25) = 1.674, *p* = .2075], increased freezing across conditioning days [main effect: *F*(2, 50) = 44.685, *p* < .0001], and displayed noise-elicited decreases in freezing during conditioning [main effect: *F*(1, 25) = 69.535, *p* < .0001]. Of note, female rats in this experiment showed slightly higher levels of freezing during habituation [main effect: *F*(1, 25) = 5.208, *p* = .0313], but no sex differences were observed across conditioning days [*F*(1, 25) = .768, *p* = .3891]. After conditioning, rats were either placed back into the conditioning context to extinguish contextual fear for 45 min (Ext) or they were exposed to the habituation context for an equal amount of time (No-Ext) (**Figure 3A**, **Day 5**). Although there was not a significant group X time interaction during extinction [*F*(1, 27) = 2.994, *p* = .095], planned comparisons revealed that Ext animals showed a significant reduction in freezing [*F*(1, 14) = 16.930, *p* = .0011], whereas No-Ext animals showed stable and lower levels of freezing during the session [*F*(1, 13) = 1.611, *p* = .2266].

**Figure 3.**
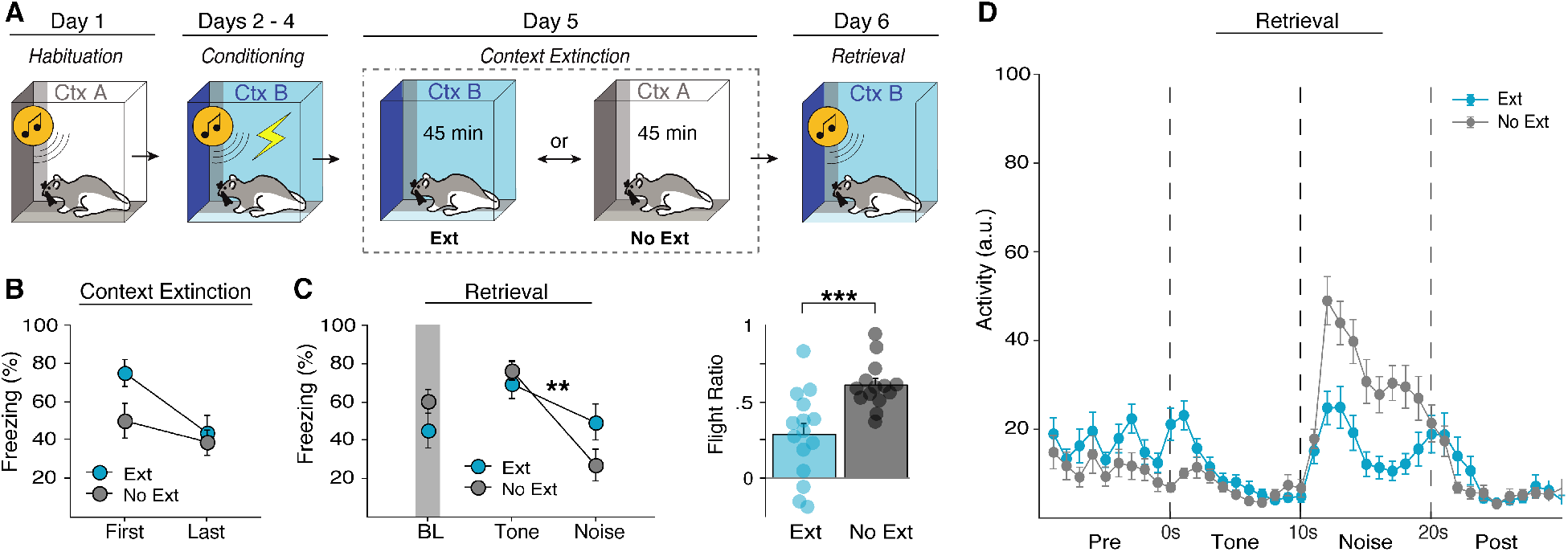
Extinguishing contextual fear reduces flight-like behavior. **A**, behavioral design for Experiment 3. **B**, Average freezing data showing that rats that underwent context extinction (Ext) froze at high levels at the beginning of extinction which reduced by the end of extinction. Rats that did not undergo extinction (No-Ext) did not show a reduction in freezing from the first to last block of context exposure. **C**, Ext animals did not show a significant reduction in baseline freezing, but did show a reduced flight response as shown by the reduced flight ratio. **D**, Averaged activity data showing that Ext animals showed reduced activity during the white noise stimulus compared to No-Ext animals. All data are represented as mean ± SEM; *, **, and *** denotes *p*<.05, *p*<.01, and *p*<.001, respectively.

For retrieval testing, all rats were placed back into the conditioning context and presented four SCS-alone trials. Although baseline freezing was similar between groups [*F*(1, 27) = 1.857, *p* = .1843], NoExt rats showed a greater reduction in freezing to noise onset relative to Ext animals [stimulus type X group interaction, *F*(1, 25) = 15.880, *p* = .0005], which was again mirrored by changes in activity [*F*(1, 25) = 6.995, *p* = .0139], and flight ratio [*F*(1, 25) = 14.212, *p* = .0009] (**Figure 3C**). In other words, context extinction reduced flight-like behavior (**Figure 3D**). There was no main effect of sex for any of these metrics during retrieval testing [*F*(1, 25) = 1.388, *p* = .2498]. These results provide converging evidence that SCS-driven flight-like behavior is driven by summation of fear to the SCS and conditioning context.

### Flight-like responses in rats are specific to white noise and not due to sensitization

One outstanding question is whether flight behavior in rats is driven by threat imminence or stimulus salience. Predatory imminence theory posits that defensive responding scales with threat proximity, and thus, white noise may elicit flight in the SCS paradigm because it is temporally proximal to the shock US. However, recent work shows that SCS-elicited flight behavior in mice is driven by stimulus salience, specifically intensity and high frequency components of white noise (Hersman et al., 2020). Indeed, this work shows in mice that flight behavior is specific to white noise regardless of whether it precedes or follows the pure tone component of the SCS. Moreover, Hersman and colleagues show that flight behavior does not require SCS-US pairings, insofar as an unpaired SCS-US procedure also produced flight to the SCS. This suggests that sensitization or pseudoconditioning might contribute to flight to the SCS. We therefore sought to determine whether the temporal order of the stimuli in the SCS influences the emergence of flight behavior, and if flight-like behavior in rats occurs after unpaired SCS-US trials. To test this (**Figure 4A**), after habituation animals underwent either a standard SCS-US procedure described thus far (Standard), a standard SCS with a 60-s delay before the US (Unpaired), or a reversed order SCS (noise-tone) immediately followed by a US (Reversed). All groups were then tested by presenting the same SCS that they were conditioned with (either standard or reversed) without US presentation in the conditioning context. To accurately reflect freeze-to-flight transitions in this experiment the flight ratio was calculated for each group as activity during white noise relative to the 10-sec period prior to white noise onset for the first retrieval test trial. Thus, for Standard and Unpaired groups the flight ratio is the same as previously described, the ratio of the difference of noise and tone load-cell activity to the sum of noise and tone load-cell activity; however, for the Reversed group this becomes the ratio of the difference of noise and pre-SCS load-cell activity to the sum of noise and pre-SCS load-cell activity.

**Figure 4.**
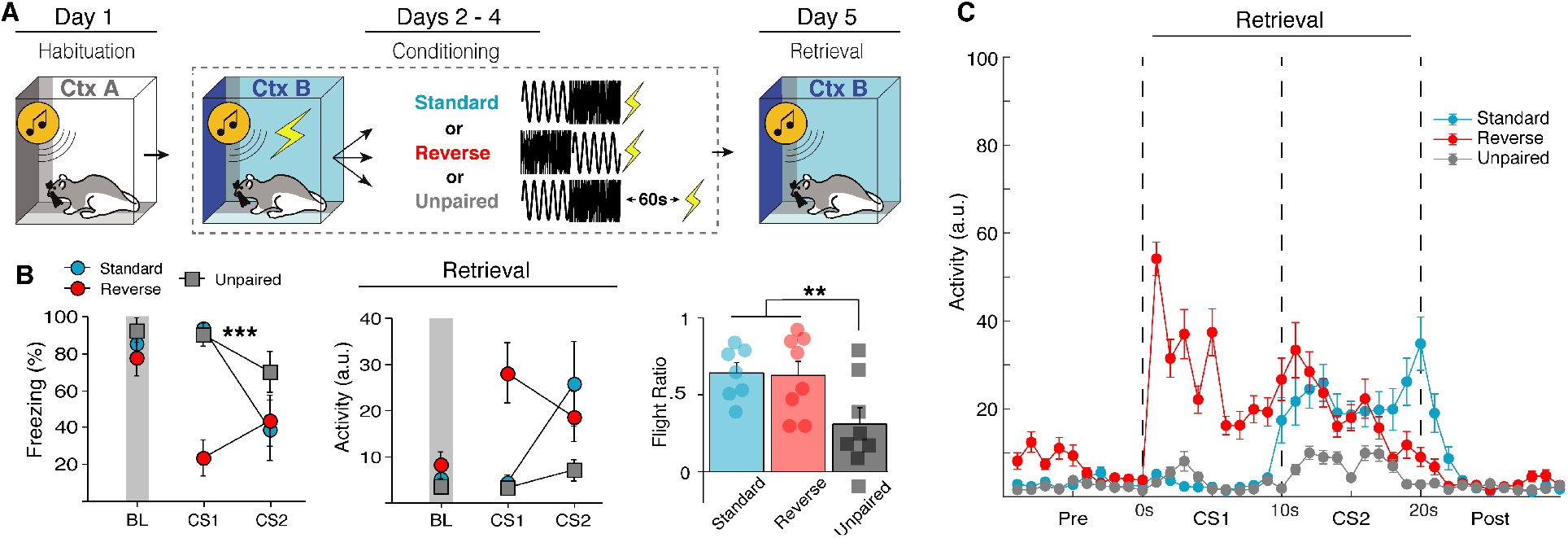
Flight-like responses in rats are specific to white noise and not due to sensitization. **A**, Behavioral design for Experiment 4. **B**, Average freezing and activity data showing that flight-like behavior is specific to the white noise stimulus regardless of the temporal of order of the SCS. In a Reversed order SCS, rats show a decrease in freezing and corresponding increase in activity to the first stimuli (noise) rather than the second (tone). The data additionally show that unpairing the SCS and US with a 60-s gap (Unpaired) prevents flight-like behavior compared to Standard SCS-US controls. This all further shown by averaged activity time across time (**C**). All data are represented as mean ± SEM; *, **, and *** denotes *p*<.05, *p*<.01, and *p*<.001, respectively.

Prior to conditioning, there was no main effect of either group [*F*(2, 17) = .858, *p* = .4416], sex [*F*(1, 17) = 2.232, *p* = .1535], or stimulus type [*F*(1, 17) = .536, *p* = .4742]; though female rats in the Reversed group did show increased freezing to tone presentations across habituation which yielded a trial X sex interaction [*F*(3, 51) = 3.777, *p* = .0160]. All animals showed increased freezing across conditioning sessions and all groups showed decreased freezing to the noise relative to the tone, including animals in which the SCS order was reversed [day X stimulus type X group interaction: *F*(4, 34) = 15.466, *p* < .0001]. This suggests that flight responses are indeed specific to white noise, rather than determined by threat proximity. Female rats in this experiment generally showed higher freezing levels across conditioning [main effect of sex: *F*(1, 17) = 5.509, *p* = .0313], although no interactions with sex were seen. During retrieval testing, Unpaired animals showed a reduced flight response to white noise [main effect: *F*(2, 17) = 7.646, *p* = .0043] compared to both Standard [*p* = .0103] and Reversed groups [*p* = .0106] (**Figure 4B**). No main effect of sex was seen [*F*(1, 17) = 1.397, *p* = .2535]. In summary, we show here that flight behavior is also exclusive to white noise in rats and suggests that footshock sensitization cannot account for flight behavior. This reaffirms previous findings reported in mice that innately aversive auditory stimuli drive flight response in the SCS paradigm, not threat imminence.

### An unconditioned SCS fails to evoke flight behavior in a threatening context

As shown in the previous experiment, flight behavior is specific to white noise and appears to be driven by stimulus salience. This raises an alternative possibility that SCS-evoked flight may be nonassociative. White noise is commonly used as an acoustic startle stimulus and it is well known that a startle response can be potentiated when presented in a shock-associated context, a process known as fear-potentiated startle (Davis and Walker, 2014; Luyten et al., 2011; McNish et al., 1997). Although we showed in the last experiment that unpaired SCS/US presentations fail to produce robust flight behavior, an unpaired CS can come to act as a conditioned inhibitor (i.e., safety signal) which may have reduced flight behavior (Rescorla, 1969). Thus, we next investigated whether an excitatory context might drive a potentiated startle to the noise that could account for SCS driven reductions in freezing and concomitant flight behavior.

All animals were first habituated with four SCS-alone trials. No differences were seen between groups [*F*(1, 24) = .604, *p* = .445], stimulus type [*F*(1, 24) = 3.205, *p* = .0860], or sex [*F*(1, 24) = .927, *p* = .4445]; although females showed slightly higher freezing across habituation trials [trial X sex interaction: *F*(3, 72) = 2.870, *p* = .0423]. To test if SCS driven flight can be explained by fear-potentiated startle (**Figure 5A**), animals either underwent standard SCS-US conditioning or received an equal number of unsignaled USs across three consecutive days. The day after conditioning, animals were placed into a novel context C where they either received five US-alone presentations or context exposure, similar to Experiment 2. Finally, all animals were presented SCS-alone trials in context C on the last day of experimentation. This creates a 2×2 design where animals were conditioned with either SCS-US or US-alone trials and were subsequently tested in either a threatening (Shock) or neutral context (No-Shock). If SCS driven flight is merely a potentiated startle response, we would expect both animals conditioned with SCS-US and US-alone trials to exhibit flight-like behavior in the shock-associated context.

**Figure 5.**
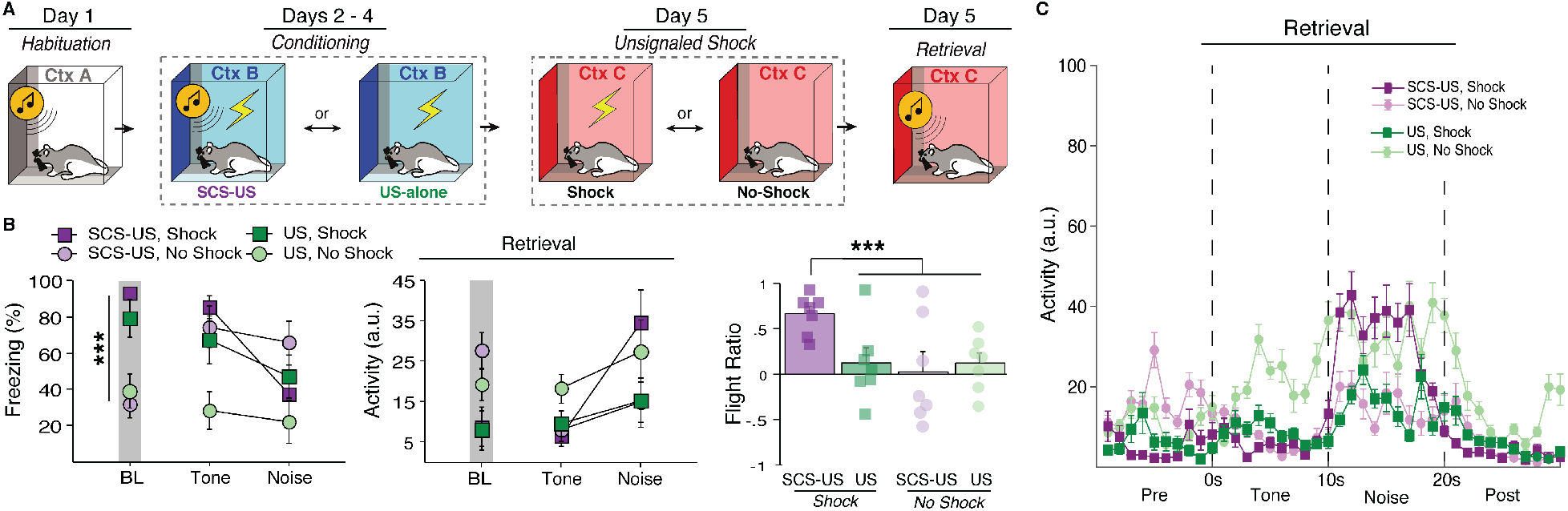
An unconditioned SCS fails to evoke flight behavior in a threatening context. **A**, Behavioral design for Experiment 5. **B**, Averaged freezing and activity data showing that rats that received US-alone trials throughout conditioning (US) and tested in a US-associated context (Shock) had a reduced flight response in comparison to rats that received SCS-US pairings. This is shown by increased freezing and decreased activity to the white noise stimulus, and a reduced flight ratio. Rats that were tested in a neutral context (No-Shock) also showed reduced flight responses compared to SCS-US/Shock animals. This is further shown by the average activity trace of each group (**D**). All data are represented as mean ± SEM; *, **, and *** denotes *p*<.05, *p*<.01, and *p*<.001, respectively.

During SCS-US conditioning, all animals showed increased freezing across sessions [*F*(2, 24) = 70.058, *p* < .0001] and lower levels of freezing to noise than tone [*F*(1, 12) = 29.132, *p* = .0002], though the difference in freezing between stimuli were similar across days [day X stimuli: *F*(2, 24) = 2.784, *p* = .0818]. Animals that received US-alone trials also showed increased freezing across sessions [*F*(2, 24) = 22.404, *p* < .0001]. No sex differences were observed during conditioning. One day after conditioning, animals either received five unsignaled footshocks in a novel context C or were exposed to the context for an equal amount of time. Animals that received footshocks showed an increase in freezing behavior relative to No-Shock animals [*F*(5, 120) = 10.762, *p* < .0001]. Although there was no main effect of sex [*F*(1, 24) = .445, *p* = .5111], there was a trial X group X sex interaction [*F*(35, 120) = 2.507, *p* = .0338], again driven by slightly higher freezing in female No-Shock animals. On the day of retrieval testing (**Figure 5B**), Shock animals showed much higher levels of baseline freezing than No-Shock animals [*F*(1, 23) = 34.997, *p* < .0001] and there was no difference in SCS-US and US-alone groups [*F*(1, 23) = .145, *p* = .7070] or sex [*F*(1, 23) = .064, *p* = .8025]. At the first SCS presentation, all groups froze at a high level except for the US-alone/No-Shock group [conditioning X unsignaled shock interaction: *F*(1, 23) = 4.062, *p* = .0557]. Upon white noise onset, the SCS-US/Shock group showed a dramatic increase in activity greater than all other groups [flight score main effect: *F*(3, 24) = 3.632, *p* = .0272], including the US-alone/Shock group [*p* = .0193] (**Figure 5B**). Indeed, the US-alone/Shock group showed similar flight-like behavior as animals that were never shocked in that context. We therefore conclude that SCS driven flight behavior does require a direct SCS-US pairing in this paradigm and thus cannot be attributed to sensitization or fear potentiated-startle.

### Muscimol inactivation of the central or extended amygdala attenuates flight behavior

If SCS-evoked flight depends on context-US associations, then inactivating brain regions that are critical for this process should block flight behavior. The bed nucleus of the stria terminalis (BNST) has been shown to be critical for the expression of contextual but not cued fear (Davis et al., 2010; Goode and Maren, 2017; Sullivan et al., 2004), whereas the CeA is critical for both contextual and cued fear (Janak and Tye, 2015; Maren and Quirk, 2004). Based on this, we reasoned that inactivation of the either CeA or BNST would block freezing to the conditioning context and SCS-driven flight responses.

All rats were implanted with cannula targeting either the BNST or CeA one week prior to SCS habituation and conditioning (**Figure 6A-C**). During habituation, there were main effects of stimulus type [*F*(1, 40) = 5.145, *p* = .029] and trial [*F*(3, 120) = 3.284, *p* = .023] with animals showing increased freezing to tones at the end of habituation. All animals showed increased freezing to the SCS across conditioning days [*F*(2, 80) = 117.994, *p* < .0001] with decreased freezing to the white noise stimulus [main effect of stimulus type: *F*(1, 40) = 356.902, *p* < .0001]. No sex differences were seen across habituation or conditioning. Immediately prior to retrieval testing rats received local infusions of either the GABAA agonist muscimol (MUS) or saline (SAL). Inactivation of either the CeA [*p =* .0013] or BNST [*p =* .0012] reduced baseline freezing relative to SAL controls [*F*(1, 40) = .901, *p* = .3482], an indication of diminished contextual fear. CeA animals showed lower freezing to the tone [main effect: *F*(2 40) = 5.457, *p* = .008] compared to both SAL [*p* = .0028] and BNST animals [*p* = .038] (**Figure 6D**). Upon noise onset, there was not a stimulus type X group interaction [*F*(1, 40) = .901, *p* = .3482], but planned comparisons reveal that SAL animals [*p* < .0001], but not CeA [*p* = .6165] or BNST [*p* = .1321], showed a reduction in freezing behavior (**Figure 6D**). This is reflected in the flight ratios [main effect: *F*(2, 40) = 8.773, *p* = .0007] that show both CeA [*p* < .0001] and BNST inactivation [*p* = .0067] blunted flight response (**Figure 6G**). This can further be seen in averaged activity plots (**Figure 6E-F**). No sex differences were seen during retrieval. In summary, inactivating either the CeA or BNST was sufficient to block both contextual fear and SCS-evoked flight responses. This provides further evidence that SCS flight is a high fear state gated by a summation of contextual and cued fear.

**Figure 6.**
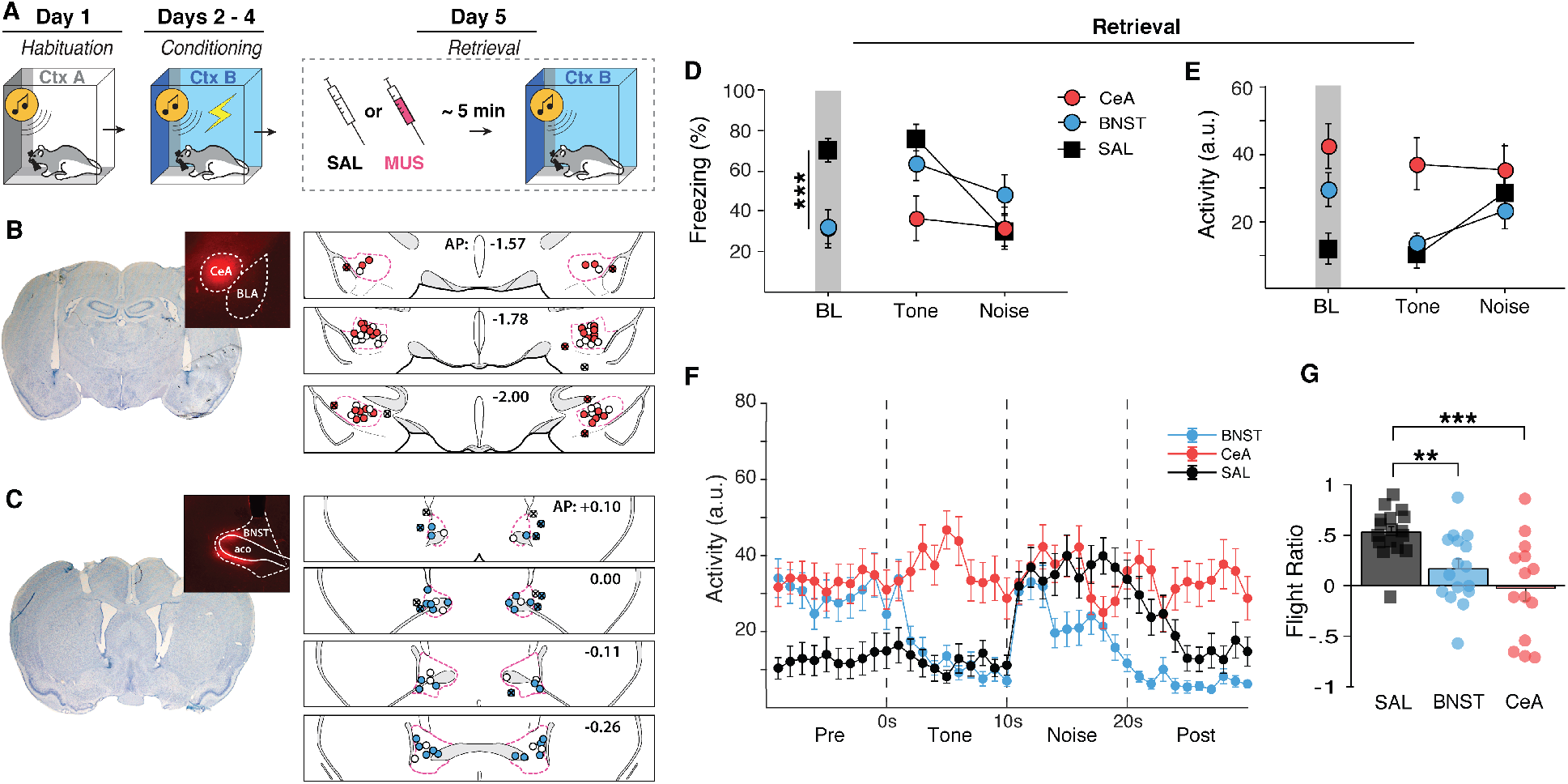
Pharmacological inactivation of either the BNST or CeA disrupts flight-like behavior. **A**, Behavioral design for Experiment 6. Histological summary of CeA (**B**) and BNST (**C)** cannula placements with representative thionin-stained sections and drug spread with fluorescent muscimol. Labeled anterior-posterior coordinates are relative to bregma. **D**, averaged freezing data showing that CeA and BNST animals both showed reduced baseline freezing. BNST animals increased freezing to tone presentation and remained at higher freezing levels during white noise. CeA animals remained at low levels of freezing during the SCS. **E** and **F**, averaged activity levels during the first SCS presentation showing that BNST animals showed a reduced flight response and CeA animals did not increase activity from tone to noise at all despite a lack of freezing behavior. This is reflected in the flight ratio (**G**). All data are represented as mean ± SEM; *, **, and *** denotes *p*<.05, *p*<.01, and *p*<.001, respectively.

## DISCUSSION

Here we investigated context-dependent flight behavior evoked by a serial-compound stimulus conditioning procedure in male and female rats. First, we show that context-dependent flight-like behavior can be evoked in rats using the SCS procedure, although jumping was less frequent than previously reported in mice (Fadok et al., 2018). We further found that flight occurs in any shock-associated context and that extinguishing contextual fear to the conditioning chamber suppresses flight-like behavior, demonstrating that SCS-evoked flight behavior reflects the summation of cued and contextual fear. Although flight is specific to white noise, we found that fear-potentiated startle and sensitization could not account for SCS-evoked flight. That is, neither unpaired nor neutral SCS presentations were sufficient to drive flight behavior in a shock-associated context, even when animals had received an equal number of prior footshocks. Finally, we show that pharmacological inactivation of brain regions that are critical for the expression of contextual fear, either the CeA or BNST, is sufficient to block the expression of flight behavior. Together, these data demonstrate that conditioned flight behavior in the SCS paradigm is driven by a high fear state via a combination of contextual and cued fear.

Until now only mice had been used to study flight responses in the SCS paradigm and it was unknown if rats would also elicit flight behavior to an SCS (Dong et al., 2019; Fadok et al., 2017; Hersman et al., 2020). We found that the SCS procedure indeed evoked context-dependent flight responses similar to reports in mice, although jumping is less frequent in rats. One important limitation of our study is that we used indirect measures of motion, opposed to other work that used direct measurements of speed via camera which makes direct comparisons of locomotion difficult (Dong et al., 2019; Fadok et al., 2017). Despite this, visual comparison of the increase in activity seen here to those in previous reports appear similar (Dong et al., 2019; Fadok et al., 2017; Hersman et al., 2020). Additionally, increases in activity and jumping seen here were both specific to the white noise stimulus. It is currently unclear why rats display infrequent jumps although we speculate that this may be a species-specific difference.

Previous work suggests that female rats display more active defensive behaviors, such as defensive darting, in response to an aversive CS (Gruene et al., 2015). Based on this we expected that female rats may be more likely to show flight-like behavior compared to males in the SCS paradigm. However, we did not observe significant sex differences in SCS-evoked jumping or activity levels. Darting behavior was seldom observed during SCS presentations in both female and male rats. This could be due to scoring differences as original reports used an automated detection method (Gruene et al., 2015). Conversely, other work shows that female mice actually exhibit increased freezing in the SCS paradigm, though the average speed of male and female mice during the SCS did not differ (Borkar et al., 2020). We do report here that female rats frequently show increased freezing during SCS habituation and conditioning, although the effects were small and not always present across experiments. Assessing divergent defensive strategies in male and female rats may require machine learning-based behavioral scoring methods (Mathis et al., 2018; Pereira et al., 2020).

As others have shown (Hersman et al., 2020), we found that rats display flight behavior specifically to white noise, even when the order of the SCS is reversed. We further show that flight is not due to sensitization and that flight cannot be explained by fear potentiated startle. This supports previous work demonstrating that stimulus salience determines flight behavior in mice (Hersman et al., 2020). This previous work specifically shows that it is the high-frequency component and intensity (>80 dBs) of white noise that evoke flight. Indeed, loud, high-frequency stimuli appear to innately produce flight behavior in mice. However, it was also shown that sensitization by previous US presentations actually reduces the frequency of flight behavior due to increased competition with freezing behavior (Mongeau et al., 2003). So, how do we reconcile that a direct SCS-US association is necessary to drive flight behavior to a white noise stimulus in rats? We believe that the most parsimonious explanation is that flight in the SCS paradigm is driven by a high fear state in which a threshold is reached such that a freeze-to-flight transition occurs. Specifically, SCS flight is driven by the summation of cued, contextual, and innate fear; although all three are not always necessary to elicit flight behavior. For example, flight in mice can be evoked innately to loud, high-frequency stimuli (i.e., without conditioned fear) (Mongeau et al., 2003). Additionally, flight to an SCS can be evoked without a salient high-frequency component by increasing the intensity above 90 dBs (Hersman et al., 2020). In the SCS paradigm, auditory stimuli are presented at 75-80 dB which appears to be just below the threshold to innately evoke flight responses to white noise (Fadok et al., 2017; Hersman et al., 2020). Coupled with our findings that SCS flight requires both SCS-US and context-US associations, we propose that cued and contextual fear act in sum with stimulus salience to cause a freeze-to-flight transition in the SCS paradigm.

In line with our behavioral results, we found that reversible inactivation of either the CeA or BNST is sufficient to disrupt not only contextual fear, but also context-dependent flight responses in the SCS paradigm. The finding that CeA inactivation disrupted both contextual and cued fear is supported by decades of work demonstrating that the CeA is critical to the expression of conditioned responses (Janak and Tye, 2015; Killcross et al., 1997; Ressler et al., 2020). Moreover, our finding that inactivating the BNST disrupted defensive freezing to the conditioning context, but not the SCS, is in line with previous work showing that the BNST mediates fear to unpredictable threats (Davis et al., 2010; Goode et al., 2019, 2015; Goode and Maren, 2017; Resstel et al., 2008; Sullivan et al., 2004; Walker et al., 2009; Zimmerman and Maren, 2011). In the original report by Fadok and colleagues, they found that SCS-elicited flight is gated by neurons in the CeA expressing corticotrophin-releasing hormone (CRH+) that inhibit somatostatin-expressing neurons (SOM+) (Fadok et al., 2017). Stimulation of SOM+ neurons can elicit freezing behavior in naïve animals (Li et al., 2013; Penzo et al., 2015, 2014; Yu et al., 2016), however, flight responses evoked by stimulation of CRH+ requires prior conditioning (Fadok et al., 2017). This raises the possibility that SCS-evoked flight behavior may be evoked indirectly via the inhibition of CeA-driven freezing behavior, and thus, may not directly require the CeA. We extend this literature by showing that the CeA is indeed required for SCS-evoked flight behavior by showing that reversible inactivation CeA attenuates flight responses. Collectively, this findings detail for this first time how conditioned auditory and contextual fear may sum with stimulus salience to drive circa-strike behavior.

Considering this, what neural structure may then be responsible for the proposed threshold-like mechanism gating flight behavior? The periaqueductal grey (PAG) is a midbrain structure that is critical to defensive responding downstream of both the CeA and BNST (Tovote et al., 2016, 2015) and appears to have a specialized role in mediating escape behavior (Lefler et al., 2020). Functional differences in the PAGs dorsal-ventral axis exists such that the ventral PAG mediates freezing behavior and the dorsal PAG mediates flight behavior (Assareh et al., 2016; Carrive, 1993; Franklin, 2019; Vianna et al., 2001). For example, stimulation of the dorsal PAG can result in both flight and freezing behavior, whereas stimulation of the ventral PAG results exclusively in freezing (Assareh et al., 2016; Carrive, 1993; Chou et al., 2018; Kim et al., 2013; Vianna et al., 2001). Importantly, recent work in mice has indeed shown that the dorsal PAG performs a synaptic threshold mechanism for computing escape behavior in a looming-disc paradigm (Evans et al., 2018). Based on this, we speculate that concurrent CeA and BNST input to the PAG could drive the threshold-like mechanism underlying context-dependent flight behavior (Nagy and Paré, 2008). Alternatively, BNST projections to the CeA may represent necessary inputs to drive CeA CRH+ neurons to gate flight behavior (Fadok et al., 2017; Gungor et al., 2015; Gungor and Paré, 2016; Yamauchi et al., 2018). Future work should investigate these pathways and their potential role in mediating flight behavior in the SCS paradigm.

To summarize, we have shown that rats display flight-behavior in the SCS paradigm similarly to mice, although rats show less frequent escape-like behaviors such as jumping and darting. Flight-like behavior evoked by the SCS is specific to white noise, gated by contextual fear, and cannot be accounted for by sensitization or fear potentiated startle. We conclude that SCS conditioning results in a high fear state driven by the summation of cued, contextual, and innate fear that drives a freeze-to-flight transition. Future work should investigate the neural mechanisms underlying the transition from post-encounter to circa-strike defensive behaviors and how this is driven by a combination of conditioned and innately aversive stimuli (Fanselow and Lester, 1988; Mobbs et al., 2020). This may reveal important clinical implications for psychiatric disorders that are characterized by high fear states and the dysregulation of contextual processing, such as panic disorder and PTSD (Goddard, 2017; Maren et al., 2013).

## MATERIALS AND METHODS

### Subjects

Experiments used adult Long-Evans rats (*n* = 163) acquired from Envigo (Indianapolis, IN; 200-240 g upon arrival). Males and females were used in equal numbers throughout all experiments. All animals were housed in a climate-controlled vivarium and kept on a fixed light/dark cycle (lights on starting at 7:00 AM and off at 9:00 PM; experiments took place during the light phase of the cycle). Rats were individually housed in clear plastic cages (with bedding consisting of wood shavings; changed weekly) on a rotating cage rack. Group assignments for behavioral testing was randomized for cage position on the racks. Animals had access to standard rodent chow and water *ad libitum*. Animals were handled by the experimenter(s) (~30 sec/day) for five consecutive days prior to the start of any surgeries or behavior. All procedures were in accordance with the US National Institutes of Health (NIH) Guide for the Care and Use of Laboratory Animals and were approved by the Texas A&M University Institutional Animal Care and Use Committee.

### Apparatuses

All behavioral testing occurred within one of two rooms in the laboratory. Each behavioral room housed eight identical rodent conditioning chambers (30 cm × 24 cm × 21 cm; MED Associates, Inc.). Each chamber was housed in a larger, external sound-attenuating cabinet. Rear walls, ceilings, and the front doors of the testing chambers were made of Plexiglas, while their sidewalls were made of aluminum. Grid floors of the chambers were comprised of nineteen stainless steel bars (4 mm in diameter) and spaced 1.5 cm apart (center to center). The grid floors were attached to an electric shock source and a solid-state grid scrambler for delivery of the US (MED Associates, Inc.). A speaker attached to each chamber was used to deliver the auditory CS. As needed for each context, the chambers were equipped with 15 W house lights, and small fans were embedded in the cabinets (providing background noise of ~70 dB). An aluminum pan was inserted beneath the grid floor to collect animal waste. A small camera was attached to the top of the cabinet for video monitoring of behavior.

Measurements of freezing and motor activity were performed using an automated system (Maren, 1998). Specifically, each behavioral testing chamber rested on a load-cell platform that was sensitive to cage displacement due to each animal’s movements. During behavioral testing, load-cell activity values (ranging from −10 to +10 V) were collected and digitized at 5 Hz using Threshold Activity Software (MED Associates, Inc.). Offline conversions of the load-cell activity values were performed to generate absolute values ranging from 0 to 100; lower values indicate minimal cage displacement, which coincided with freezing behaviors in the chambers. Accordingly, freezing bouts were defined as absolute values of ≤10 for 1 s or more. The percentage of freezing behavior during the pre-SCS baseline and SCS trials was computed for each behavioral session. Motor activity was analyzed by directly reporting the absolute values generated by the Threshold Activity Software (i.e., larger values indicated more movement in the cage). Jumping and darting behavior were manually scored off-line from video recordings by an experimenter blind to experimental conditionings.

Unique contexts (A, B, and C) were used for various phases of behavior testing. Chamber assignments were unique to each context and group assignments were counterbalanced across test chambers when possible. For each experiment contexts A and B were assigned to different behavioral testing rooms. For context A, the test chamber and pans beneath the grid floors were wiped down with an ammonium hydroxide solution (1%). The cage lights were turned off, chamber fans were turned on, and the cabinet doors were left open. Black Plexiglas panels were also placed over the grid floors. The behavioral room was lit with white light (red lights were turned off). Animals were transported to and from the chambers using white plastic transport boxes. For context B, an acetic acid solution (3%) was used to wipe down and scent the chambers, the cage lights were turned on, the chamber fans were turned off, and the cabinet doors were closed. The behavioral room was lit with dim red light (white room lights were turned off). Rats were transported to and from context B using black plastic transport boxes that included a layer of clean bedding. For context C, an ethanol solution (70%) was used to wipe down and scent the chambers, the cage lights were turned on, the chamber fans were turned on, and the cabinet doors were open. The behavioral room was lit with white lights (red lights remained off) and rats were transported to and from context C in white plastic transport boxes with clean bedding. Testing in context C was always performed in the same behavioral room as context A.

### Experimental Design

Overviews of each behavioral experiment are provided in the figures. The auditory serial-compound stimulus (SCS) used for all experiments was comprised of ten 500 ms pure tone pips (80 dB, 7 kHz) presented at a frequency of 1 Hz (500 ms inter-pip intervals, 10 s total length) and immediately followed by ten 500 ms white noise pips (80 dB, 1-20 kHz) presented at a frequency of 1 Hz (500 ms inter-pip intervals, 10-s total length). During conditioning the SCS was paired with a mild unconditioned footshock stimulus (US, 1.0 mA, 2 sec), unless noted otherwise. Intertrial-intervals (ITI) were always 60 seconds.

#### Experiment 1: Flight behavior in the conditioning vs habituation contexts

In this experiment we tested if the SCS paradigm could evoke flight-like behavior in rats similar to what has previously been reported in mice. To this end, behavioral testing consisted of a habituation session, three conditioning sessions, and a retrieval session. Habituation sessions consisted of a 3 min pre-SCS baseline period followed by 4 SCS presentations without footshock in context A. Next, all rats underwent conditioning consisting of a 3 min pre-SCS baseline period followed by 5 SCS-US presentations in context B for three consecutive days. Multiple days of conditioning were necessary to drive flight responses (Fadok et al., 2017). Retrieval testing consisted of a 3-min pre-SCS baseline period followed by 4 SCS-alone presentations in either context A (habituation) or context B (conditioning). A within-subject design was used such that half the rats were tested in either the habituation or conditioning context first and later were tested in either the conditioning or habituation context, respectively, the subsequent day (counter-balanced for test order). There was only one group tested in this experiment using a within-subjects design (*n* = 8). No animals were excluded.

#### Experiment 2: Unsignaled footshock

For this experiment we wanted to determine if flight-like behavior could be driven in other shock-associated contexts. Behavioral testing for this experiment consisted of a habituation session (context A), three conditioning sessions (context B), a session where rats either received unsignaled footshocks (context C), and a final between-subjects retrieval session in context C. Habituation and conditioning sessions were identical to Experiment 1. For the unsignaled footshock session, all rats were placed into a novel context C where half were presented a 3-min pre-stimulus baseline followed by 5 unsignaled footshocks (1.0 mA, 2 sec) with 60-sec ITIs (Shock), and the other half were merely exposed to the context for an equal amount of time (No-Shock). The next day all rats were returned to context C where they were presented with 1 SCS-alone trial following a 3 min baseline period. The groups were as follows: Shock (*n* = 8) and No-Shock (*n* = 8). No animals were excluded.

#### Experiment 3: Context extinction

In this experiment we tested if flight behavior could be diminished by extinguishing contextual fear. Behavioral testing consisted of habituation, three days of conditioning, a context extinction session, and a retrieval session. Habituation and conditioning were identical to previous experiments. For context extinction, half of the rats underwent context extinction by exposing them to the conditioning context (context B) for 45 min for two consecutive days (EXT) while the other half were re-exposed to context A for an equal amount of time (No-EXT). The subsequent day all animals were placed back into context B where they underwent a retrieval session consisting of a 3-min pre-SCS baseline followed by 4 SCS-alone presentations. Groups were as follows: EXT (*n* = 14) and No-Ext (*n* = 15). Two EXT (*n* = 2) and one No-EXT (*n* = 1) animals were excluded as statistical outliers (flight ratio >2 SDs ± mean).

#### Experiment 4: Reverse and unpaired SCS

This experiment tested both the temporal order of the SCS and the SCS-US relationship on flight behavioral. Behavioral testing for this experiment was similar to previous experiments in that it consisted of one day of habituation (context A), three days of conditioning (context B), and a retrieval session (context C). However, conditioning consisted of either 5 standard SCS-US pairings (Standard), 5 reversed (noise followed by tone) SCS-US pairings (Reverse), or 5 unpaired presentations with a 60-sec delay between each SCS and US (Unpaired). Following the final day of conditioning, all rats were placed back into context B and presented 4 SCS-alone presentations. Groups are as follows: Standard (*n* = 8), Reverse (*n* = 8), and Unpaired (*n* = 8). No animals were excluded.

#### Experiment 5: Contextual fear potentiation

For this experiment we wanted to determine if the context dependence of flight behavior could be accounted for by fear potentiated startle. This experiment is designed similar to Experiment 2 which consisted of habituation (context A), three days of conditioning (context B), unsignaled footshock (context C), and retrieval (context C). However, following habituation, rats were either conditioned with 5 SCS-US pairings (SCS-US) or received 5 unsignaled footshocks (US-alone) in context B for three consecutive days. After conditioning, each of these groups were placed in a novel context C where half received unsignaled footshocks (Shock) and half were merely exposed to the context for the same amount of time (No-Shock), thus creating a 2×2 design. On the final day, all rats were placed in context C and presented 10 SCS-alone trials after a 3-min baseline. Groups were as follows: SCS-US/Shock (*n* = 8), SCS-US/No-Shock (*n* = 8), US-alone/Shock (*n* = 8), and US-alone/No-Shock (*n* = 8). No animals were excluded.

#### Experiment 6: Muscimol inactivation of the CeA and BNST

In Experiment 6 we tested if flight behavior that depended on contextual fear could be blocked by inactivating regions necessary for the expression of context fear. Rats were first chronically implanted with bilateral cannula targeting either the CeA or BNST. One week after surgery, all rats underwent habituation (context A) and three days of conditioning (context B) as previously described. Immediately prior to retrieval testing, rats received 0.3-μl microinfusions of either the GABAA agonist muscimol (0.1 μg /μl) or saline at a rate of 0.3 μl/min and infusions needles stayed in-place for at least 2 minutes post-infusion. They were then placed into transport boxes and moved to the behavioral testing room for retrieval testing took place as previously described with a 3-min baseline period and 4 SCS-alone trials. Groups were as follows: CeA (*n* = 14), BNST (*n* = 14), SAL (*n* = 16).

### Surgery

Rats were anesthetized with isoflurane (5% induction, ~2% maintenance), the top of their heads were shaven, and they were placed in a stereotaxic mount (Kopf Instruments, Tujunga, CA). A small incision was made with a scalpel, fascia lining the skull was scrubbed away with cotton swabs, and the scalp was retracted with forceps. The skull was leveled horizontally before burr holes were drilled above either the BNST or CeA. Four additional holes were made anteriorly and posteriorly (two each) for skull screws. After skull screws were placed, two stainless-steel cannulas (26 gauge, 8mm; Plastics One) were lowered into either the CeA (target coordinates; ML: 4.0, AP: −2.0, DV: −8.0) or the BNST (target coordinates; ML: 1.5, AP: 0.0, DV: −6.5). Cannula targeting the BNST were inserted at a 10° angle to avoid rupturing the ventricle. Thus, angled coordinates used during stereotaxic surgery targeting the BNST were as follows: ML: 3.13, AP: 0.0, DV: −6.19 (ML: medial-lateral, AP: anterior-posterior, DV: dorsal-ventral). All coordinates are in reference to the skull surface at bregma. Cannula were then affixed to the skull with dental acrylic and a stainless-steel dummy (30 gauge, 9 mm; Plastics One) was inserted into the guide cannula. Rats were allowed to recover for ~1 week after surgery before behavioral testing.

### Drug microinfusions

The day of retrieval testing rats were placed into 5-gallon white buckets and moved into a room adjacent to the vivarium for microinfusions. Dummy cannula internals were removed and a stainless-steel injector (33 gauge, 9mm; Plastics One) connected to polyethylene tubing was inserted into the guide cannula. Polyethylene tubing was connected to 10-μl Hamilton syringes that were mounted in an infusion pump (Kd Scientific). Muscimol was diluted to a concentration of 0.1 μg/μl in sterile saline. Infusions were made a rate of 0.3 μl/min for 1 min and the injectors were left in place for 2 min post-infusion to allow for adequate diffusion. Each infusion was verified by movement of an air bubble that separated the drug or sterile saline from distilled water within the polyethylene tubing. Clean dummy internals were inserted into each guide cannula after infusions. All infusions were made ~5 min prior to behavioral testing.

### Histology

Twenty-four hours after retrieval testing animals were sacrificed to confirm cannula placement. Animals were overdosed with sodium pentobarbital (Fatal Plus, 100 mg/ml, 0.7 ml), transcardially perfused with ice-cold saline and fixed with 10% physiological formalin. Perfused brains were placed in physiological formalin for 14-24 hours before being moved to a 30% sucrose solution for a minimum of three days. After three days, or until brains had sunk in 30% sucrose, all brains were frozen and sectioned at −20° on a cryostat at a thickness of 40 μm. Sections were mounted onto gelatin subbed slides, thionin stained (0.25%) to better visualize cannula placement, cover-slipped with Permount (Fisher Scientific), and then imaged on a wide-field stereoscope.

A subset of animals infused with fluorescent muscimol to verify drug spread. These animals were overdosed with sodium pentobarbital and infused with 0.3 μl of fluorescent muscimol (BODIPY TMR-X conjugate; Thermo Fisher Scientific) at a rate of 0.3 μl/min. A rest period of 2 minutes was given post-infusion and then animals were immediately sacrificed. Non-perfused brains were placed in a physiological formalin solution for 14-24 hours before being placed in a 30% sucrose-formalin solution for a minimum of three days. These brains were also sectioned at −20° on a cryostat at a thickness of 40 μm. Sections were then mounted onto subbed slides, coverslipped with fluoromount (Diagnostic Bio-systems), and imaged on a fluorescent microscope at 10x resolution. Hits were confirmed by verifying that the tip of the infusion needles was within the CeA or BNST. Only animals that had bilaterally confirmed placements were included in statistical analyses. Thus, animals in which the tip of either one or both cannulas were outside of the CeA or BNST were excluded from analyses.

### Statistical analyses

All freezing and raw threshold data were analyzed offline by custom written Python and MATLAB scripts before eventual statistical testing in Statview software. All data were submitted to repeated or factorial analysis of variance (ANOVA) as described for each experiment. Fisher’s protected least significant difference (PLSD) test was used for *post hoc* comparisons of group means following a significant omnibus *F* ratio in the ANOVA (α was set at 0.05). No statistical methods were used to predetermine group sizes (group sizes were selected based on prior work and what is common for the field). Sex was included as a biological variable for all statistical comparisons. Data distributions were assumed to be normal, but these were not formally tested. All data are represented as means ± S.E.M.

## ACKNOWLEDGEMENTS

Supported by grants from the National Institutes of Health (R01MH065961 and R01MH117852 to S.M.).

## ADDITIONAL INFORMATION

### Competing interests

The authors declare no competing interests.

### Author contributions

M.T. and S.M. designed experiments; M.T., N.W., I.H., K.R., R.R., and C.O. performed the experiments; M.T. and N.W. analyzed data; M.T. and S.M. wrote the manuscript.

## Notes

### Competing Interest Statement

The authors have declared no competing interest.

